# AlphaCross-XL: a seamless tool for automated and proteome-scale mapping of crosslinked peptides onto three-dimensional protein structures

**DOI:** 10.1101/2025.01.21.633663

**Authors:** Sanjyot Vinayak Shenoy, Deeptarup Biswas, Kamal Mandal, Arthur Zalevsky, Audrey Kishishita, Ayushi Verma, Ishan Upadhay, Yi He, Andrej Sali, Rosa Viner, Sanjeeva Srivastava, Arun P. Wiita

## Abstract

Crosslinking mass spectrometry (XL-MS) is an exciting proteomics technology to capture native protein conformations in real time within biological systems. Historically, however, implementation of this technology has typically been limited to single purified recombinant proteins or *in vitro* assembled protein complexes. These limitations are associated with inherent challenges in XL-MS analysis, including extremely low abundance of crosslinked (XL) peptides and complex deconvolution of XL peptide-derived spectral data. However, impressive recent developments in computation and instrumentation have now made it feasible to address biological questions using proteome-wide XL-MS analysis. Although some XL mapping software tools exist, these require manual input of specific Protein Data Bank (PDB) structures at the single protein level, and do not function at the high throughput scale required to analyze datasets derived from thousands of proteins. To address this need, we therefore sought to develop a strategy enabling automated mapping of XL peptides onto the three-dimensional (3D) structures of proteins, at a proteome-wide scale. Herein we describe AlphaCross-XL, a first-in-class seamless computational tool for automated mapping of XL peptides onto the protein structures for intra-protein crosslinks and loop-links. The AlphaCross-XL software first retrieves protein structures from the AlphaFold Protein Structure Database and maps all the identified crosslinks onto the 3D structure. It also calculates the Euclidian distance between the crosslinked residues and reports the violated and satisfied crosslink distances based on a user-defined distance threshold, which is visually discriminated by color in PyMOL. Lastly, the tool also supports further validation of user-submitted protein structures, that can include any computer-predicted protein structure and experimentally derived protein structures (i.e., from PDB). AlphaCross-XL is available at https://github.com/sanjyotshenoy/alphacross-xl.

## INTRODUCTION

Crosslinking mass spectrometry (XL-MS) has evolved as a promising proteomics technology which can, in principle, enable proteome-scale structural profiling^1,2^. This technology relies on mass spectrometry-based determination of three-dimensional structure of proteins, facilitated by covalently locking amino acids in close proximity with chemical crosslinkers^3^. Experimentally, this is accomplished by applying to the biological sample small molecule crosslinkers which contain amino acid reactive groups connected by a spacer arm. This crosslinking is followed by regular proteolytic digestion (i.e, trypsin) and detection of the crosslinked peptides, followed by comparison to the known or predicted protein structure^4^. Lysine-targeting crosslinkers are most commonly used, given this residue’s high abundance on the exposed surface of the proteins and its efficient reactivity against NHS ester-based chemical crosslinkers such as DSSO and PhoX^5,6^. In principle, though, any amino acid can be targeted for XL-MS analysis, for example using recently described photoactivatable diazirine containing crosslinkers, which promiscuously crosslink proximal amino acids^7,8^. Regardless of crosslinker used, owing to the inherent challenges associated with this technology, including extremely low abundance of crosslinked peptides and complexities of computationally deconvolving mass spectra, the implementation of XL-MS to address biological questions has in practice primarily been limited to single protein or protein complex level structural analysis. However, proteome-scale applications of XL-MS are now more recently emerging as a powerful tool to address biological questions^9–11^. One such application we recently reported is identification of protein conformation-specific cancer immunotherapy targets^1^.

Although XL-MS provides relatively low-resolution structural information for proteins, a major advantage of XL-MS is that it can potentially inform proteome scale structural profiling of hundreds to thousands of proteins simultaneously within a cellular milieu^10,11^. This capability is in contrast to conventional biophysical methods such as crystallography, NMR, or cryogenic electron microscopy (cryo-EM), which, while high resolution, are restricted to *in vitro* artificial systems at the single protein or protein complex level^12^. However, in the current state-of-the-art for proteome-wide XL-MS, a major bottleneck for analysis relevant to protein structure involves manual interrogation of the identified crosslinked peptides by mapping them onto the known individual 3D protein structures. These structures are usually derived from repositories such as the Protein Data Bank (PDB). However, manual mapping is not feasible for proteome-scale XL-MS datasets, as there are typically hundreds to thousands of proteins with thousands of total crosslinks to map. Furthermore, many proteins of interest do not have published structures in the PDB. This gap necessitates development of computational tools for automated and proteome scale mapping of crosslinked peptides onto the 3D structure of a large dataset of proteins.

To this end, herein we describe a standalone computational tool for proteome scale mapping and visualization of XL-MS data called “AlphaCross-XL”. This tool automatically retrieves the protein structures from the AlphaFold Protein Structure Database^13^ and maps all the respective intra-protein crosslinks onto those protein structures. The input file of AlphaCross-XL contains the cross-linked peptides information in CSV format and is compatible with the search output of a variety of existing XL-MS analysis suites such as pLink 2.0^14^, Kojak^15^, SCOUT^16^, and XiSearch^17^. The output of AlphaCross-XL is a directory of proteins with their individual and consolidated mapped crosslinks onto 3D structures displayed as PyMOL session PSE files to facilitate further interrogation of cross-linked peptides for structural inferences. The tool allows color-based visual discrimination of crosslinks based on the radius of reactivity of the crosslinker used. Considering that the protein structures obtained from AlphaFold Protein Structure Database are *in silico* generated^18^, we underscore the confidence level of the mapped crosslinks based on a per-residue model confidence score (pLDDT)^18^. AlphaCross-XL also allows mapping of cross-linked peptides on selected PDB structures. This enables focused interrogation of the candidate proteins triaged out of the AlphaFold based structural mapping for further validation of conformational alterations. Such candidate proteins can then be taken forward for conformation specific biological implications in various disease contexts. The tool also has a graphical user interface (GUI) designed for ease of use even without significant computational expertise. AlphaCross-XL is hosted on Google Colab, a free browser-based integrated development environment (IDE) that offers its seamless use. Additionally, it is also available in command line version.

## EXPERIMENTAL PROCEDURES

### XL-MS Datasets Analyzed

Structural surfaceomics datasets from myeloma cell line model AMO-1 (PXD059495) and acute myeloid leukemia (AML) cell line model Nomo-1 (PXD035591) were accessed on 07-31-2024 for secondary analysis with AlphaCross-XL. Processed data files including both PhoX- and DSSO-identified crosslinks from the Nomo-1 model were downloaded from ref.^1^. For validation of AlphaCross-XL in bacterial systems, we re-analyzed processed data from Ruwolt *et al*., which contained PhoX crosslinked *E. coli* DH5a, where crosslinks were identified with Proteome Discoverer 3.0 software (Thermo Fisher Scientific) with Xlinkx node 3.0, and the CSV output of the search software was used as the input for AlphaCross-XL^19^. For the myeloma dataset, we processed DSSO-labeled datasets acquired at the MS^3^ level on Thermo Fisher Scientific Orbitrap tribrids mass spectrometers. Raw LC-MS files were analyzed with Proteome Discoverer 3.0 (Thermo Fisher Scientific) with Xlinkx node 3.0 using the cleavable search algorithm setting for crosslinked peptides and SEQUEST HT for loop links, mono-links, and regular peptides. Precursor and fragment tolerances were 20 ppm and 0.02 Da, respectively. Dynamic modifications for crosslinked peptides were set as DSSO, methionine oxidation, and protein N-term acetylation, while the fixed modification was set to cysteine carbamidomethylation. For searching, a cell line custom surfaceome database generated in-house was used^1^. The enzyme was set to Trypsin with up to 3 missed cleavages allowed. Crosslinks were filtered at CLSM level FDR < 1% separately for intra- and interlinks with 20 ppm tolerance. The filtered crosslinked result file from Proteome Discoverer was directly used as the input file for AlphaCross-XL.

### Development and Distribution of Alpha Cross-XL software

AlphaCross-XL is written in Python and primarily implemented as an IPython Notebook, which features an interactive GUI, but can also be used via command-line. To provide the research community with an easy means of analyzing and visualizing the XL-MS data we have hosted AlphaCross-XL on Google Colaboratory (or Colab) as an online version (**Fig. 1**). The primary advantage of this online version is its intuitive, user-friendly and clutter-free nature, as users need not install anything on their own devices, as the required software package dependencies are retained in the cloud. However, the use of the online interface is not required as AlphaCross-XL is also offered as an open-source software that can be installed locally and run at the command line. AlphaCross-XL currently supports intra-protein crosslinks and loop-link as the crosslink types. This tool (which runs on Python (v3.10 and above)) leverages multiple popular Python packages, including Jupyter Widgets (ipywidgets; v7.7.1; for creating interactive GUI widgets), ProDy^20^ (2.4.1; used for structural data computation), Pyfastx^21^ (2.1.0; used for parsing FASTA files), PyMOL^22^Open Source (2.6.0a0; for protein structure and crosslink visualization), Pandas^23^ (v2.0.3; for dealing with tabular data), Matplotlib (3.7.1; for statistical plotting) and Seaborn (0.13.1; for statistical plotting).

**Figure 1.**
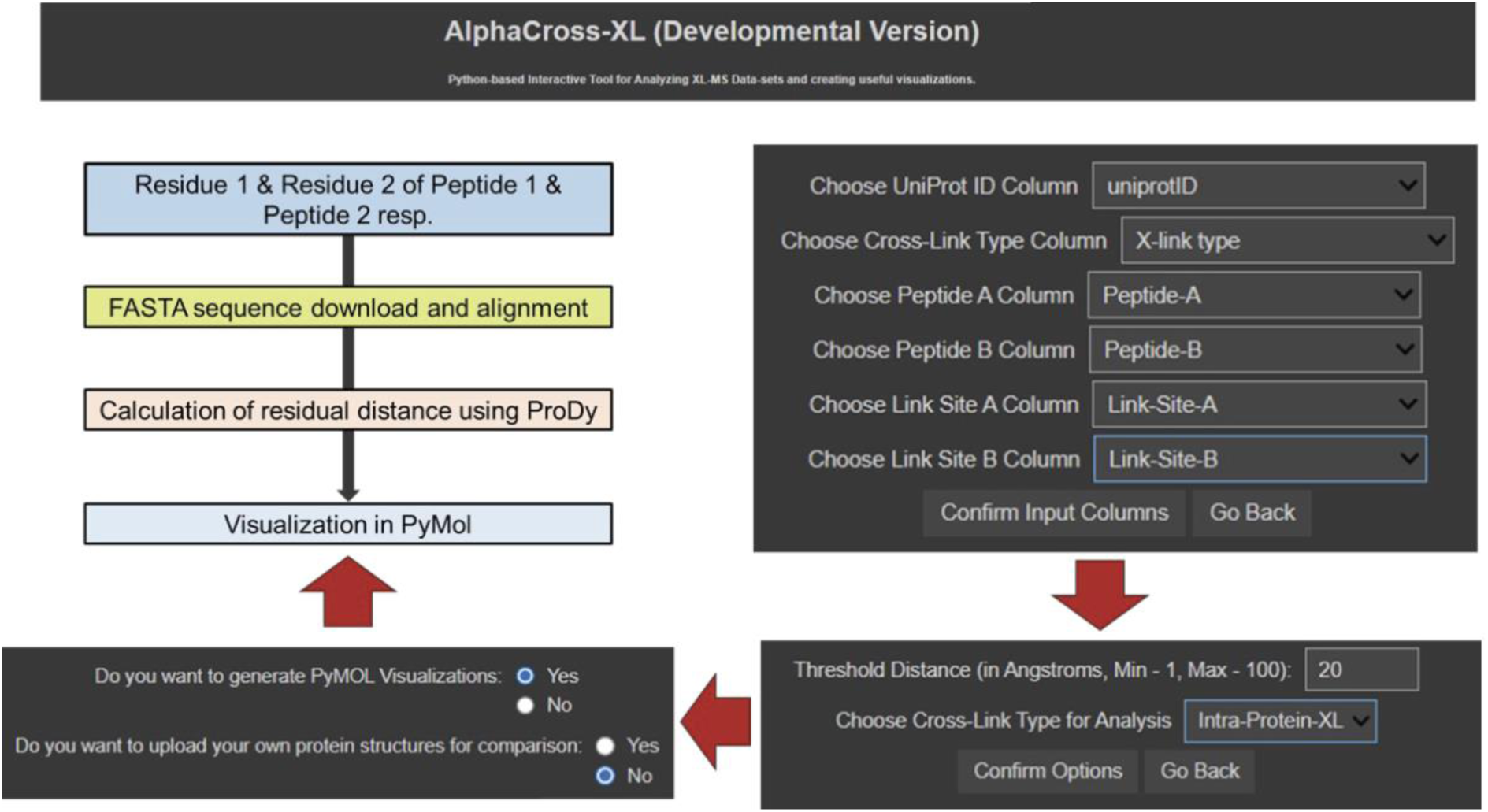
Overview of AlphaCross-XL functions and input parameters. A schematic workflow on the *left* (black arrows) is an overview of the important steps in AlphaCross-XL, including first mapping FASTA sequence acquired from UniProt, followed by mapping protein structures from the AlphaFold databases, distance calculation from ProDy, and ultimately, visualization via PyMOL. A schematic workflow starting on the *right* (red arrows) shows the user-defined input parameters required to run AlphaCross-XL, which includes assigning column types from the input CSV file, setting the crosslink distance threshold (in Ångstroms), and choosing to automatically download the generated PyMOL visualizations and/or uploading a user provided protein structure in PDB or CIF format.

### Design of AlphaCross-XL’s processing algorithm and user experience

The automation of the XL-MS analysis pipeline using AlphaCross-XL software is designed with user convenience and robustness in mind. The software’s algorithm contains a series of systematic steps for the efficient execution of XL-MS analysis, with a particular focus on performance. The tool is robust enough to deal with XL-MS data files which have different naming conventions than the standard. Subsequently, it delineates the sequential steps required to initiate XL-MS analysis, encompassing data pre-processing, retrieving the protein structure files from the AlphaFold Protein Structure Database, distance calculations, identification of violated crosslinks based on user-defined criteria, and finally creation of intuitive visualizations. Additionally, it guides users on the representation and interpretation of the output data generated by the AlphaCross-XL software. The IPython Notebook version of the tool (including the Google Colab version) has a GUI which further improves the user experience.

### Essential Input Files required by AlphaCross-XL software

To perform an analysis with AlphaCross-XL, two essential files are required: a CSV or Excel file of XL-MS search results from analysis software such as pLink 2.0^14^, Kojak^15^, SCOUT^16^, and XiSearch^17^, as well as the protein sequence database file in regular or compressed FASTA format (**Supplementary Fig. 1A**). The importance of certain columns in initiating the analysis is underscored, with emphasis placed on the necessity for the input file to contain following columns: (1) a column containing UniProt accession IDs of the protein, (2) a column designating the type of crosslink (either intra-protein crosslink or loop-link), (3) the amino-acid sequence of XL peptides in separate columns (i.e., Peptide A and Peptide B column from Kojak), (4) along with the respective crosslinking residue sites (i.e., Link-Site-A and Link-Site-B) in separate columns in peptide-centric format, meaning the format should be ‘{AA}{n}’, where ‘AA’ denotes the name of the residue in one-letter amino-acid name specification and ‘n’ denotes the relative position of the residue with respect to the first residue of the XL peptide (for example, if the peptide is LKHRK, with C-terminal Lysine residue representing the crosslink, then the link site column corresponding to this peptide should denote ‘K5’) (**Supplementary Fig. 1B)**. Peptide-centric formats are enforced as AlphaCross-XL maps the crosslinks using the XL peptides as a point of reference, enabling greater accuracy. Although AlphaCross-XL allows data files in Microsoft Excel (.xlsx) format, we recommend the input data file to be uploaded as CSV format. It is further recommended to have the UniProt reviewed sequences in the FASTA sequence database file. Since uncompressed FASTA sequence databases are large in size, we recommend uploading as a GZIP-compressed file. Users must ensure that all the sequences of all UniProt IDs mentioned in the input data file are present in the FASTA Database.

### Configuring XL-MS data analysis parameters

Once the input files have been chosen, the tool asks users to configure various analysis parameters in a sequential fashion: a) the tool will ask the user to choose the columns corresponding to the required XL-MS data attributes mentioned in “Essential Input Files required by AlphaCross-XL software”, b) after columns have been chosen, the user will be prompted to input the threshold distance (in Å) for ascertaining crosslink distance violation and choose the crosslink type for analysis (only one crosslink type can be analyzed in a single analysis run), and finally c) the user is prompted to choose whether crosslinks need to be visualized as PyMOL session files and, if so, to choose an optional ZIP archive containing appropriately-named CIF protein structure files, which will be used for comparative analysis (**Supplementary Fig. 1C)**. This ZIP archive will be extracted by the software and stored in a directory named ‘Uploaded Structures’. If the user chooses to upload their own structure files and has allowed visualization, then the visualization of crosslinks will also be performed on the user-chosen structures.

The protein structures chosen for comparative analysis are required to contain both the XL peptides with the accurate amino-acid sequence (identity match, i.e. without mutations). The structures are also required to have only one chain. This chain limit is imposed so as to prevent inter-mixing of identical, duplicate XL peptide pairs on distinct chains, usually found in homo-oligomers. In case of oligomeric proteins, the structure should be edited so as to contain both the XL peptides on a single chain.

### Crosslinking residue absolute chain position calculation for peptide-centric format

After all analysis parameters have been configured, the processing pipeline begins. First, the calculation of start positions of each XL peptide sequence’s location in the protein sequence (residue number of the peptide’s initial residue, relative to the entire protein sequence) is executed using the information found in the input data file and referred to as ‘absolute chain positions’ of the XL peptides. Subsequently, the chain position of each crosslinking residue in their respective protein’s sequence (residue number of each crosslinking residue in the entire protein sequence) is ascertained using the relative position present in link-site columns and referred to as ‘absolute chain positions’ of the crosslinking residues. These absolute positions are then used in the remainder of the analysis.

### Retrieving structure files from the AlphaFold Protein Structure Database

The AlphaFold Protein Structure Database (https://alphafold.ebi.ac.uk/) is utilized to retrieve the structure file of each unique protein (using its UniProt ID) in the input data file. The latest CIF protein structure files are fetched in real-time using AlphaFold Protein Structure Database’s API, with each protein’s structure file being stored in separate sub-directories, each named by their respective UniProt IDs, and created inside an ‘AlphaFold Structures’ directory. These files are employed to calculate the crosslinking residue distance. Subsequently, they may be used to visualize the crosslinks through PyMOL, based on user preference.

### pLDDT extraction and crosslinking residue distance calculation

The calculation of crosslinking residue distance or simply residue distance, defined as the distance between the C_a_ atoms of the crosslinked residues, is performed. To accomplish this, the ProDy package is employed. ProDy makes use of the absolute chain positions of the crosslinked residues, calculated earlier. ProDy is also utilized to extract the pLDDT score of each crosslinked residue in this step. The automation facilitates the identification of violated crosslinks distances based on the user-defined threshold C_a_-C_a_ distance dependent on the specific crosslinker used.

### Generation of PyMOL Session Files for crosslink visualization

If the user has opted for visualization of the crosslinks, AlphaCross-XL uses PyMOL’s modules to generate PyMOL session files each containing a visualized crosslink. The software creates a ‘PyMOL Sessions’ directory which contains separate sub-directories for each unique UniProt ID found in the data file. Inside a particular sub-directory of a UniProt ID, each unique crosslink found in the input data file is visualized and stored as a PyMOL Session File (PSE File). A unique crosslink for a particular protein is defined as a crosslink with a unique unordered pair of crosslinking residues with each residue identified by its respective absolute chain position (for example, if (285, 292) is a unique crosslink, then (285, 312) is unique, whereas (292, 285) is not). The PSE files are named in the format ‘{UniProt ID}-{n}’, where ‘UniProt ID’ is the UniProt ID of structure’s corresponding protein and ‘n’ is the serial number of the unique crosslink (starting with 0). Additionally, each sub-directory will contain a ‘{UniProt ID}-Consolidated’ PSE file which contains all unique crosslinks for the protein with respective UniProt ID in the same file. If the user has submitted their own structures, the crosslinks found on these structures will also be visualized and stored under the same naming convention as AlphaFold structures, but will be stored under a ‘PyMOL Sessions’ sub-directory created inside the ‘Uploaded Structures’ directory.

### Generation of Statistical Plots and Output Data Files

To provide the users with a summary of the analysis for the given input data file, AlphaCross-XL employs Matplotlib and Seaborn to generate two plots: a) A bar plot depicting all unique crosslinks on the x-axis and their residue distances on the y-axis, along with a depiction of the user-inputted threshold distance, and b) A histogram plot depicting the distribution of residue distances, with the color of the bar depicting the violation status of the frequency class. The resolution of generated plots is 300 dpi and is stored in the working directory of the tool.

After the plots have been generated, the software generates the following files – a) “acxl-alphafold-structures.zip” which is a ZIP archive containing all AlphaFold Structures retrieved and used for distance computation, b) “acxl-alphafold-pymol-sessions.zip”, which is a ZIP archive containing PSE files for AlphaFold Structures and generated only if the user has opted for visualization of the crosslinks (this ZIP archive is the compressed version of the ‘PyMOL Sessions’ directory), c) “acxl-uploaded_structs-data.zip”, which is a ZIP archive containing the protein structures submitted by the user and generated only if the user has submitted their own structures (this ZIP archive is the compressed version of the ‘Uploaded Structures’ directory, and may additionally contain visualization files under ‘PyMOL Sessions’ sub-directory if the user had opted for visualization), d) a CSV file ending with ‘_XLMS_Distances_WO_Duplicates’ containing unique rows found in the data file, each corresponding to a unique crosslink defined previously, e) a CSV file ending with ‘_XLMS_Final_Output’ containing the final output produced by AlphaCross-XL.

The web version of the software facilitates automated downloading of a ZIP archive containing all the aforementioned files generated by AlphaCross-XL and all input files used for the analysis. The local version of the software stores all the aforementioned result files in the working directory of the software.

### Curation of experimentally derived crystallographic structures for validation

To validate crosslinking data calculated from AlphaFold-derived structures, a subset of representative experimentally derived structures was also obtained from the Protein Data Bank (PDB)^24^. We focused on structures determined by X-ray crystallography and structures were then edited using BIOVIA Discovery Studio (v21.1.0.20298) to remove water molecules and hetero-atoms. If the structure had multiple chains/sub-units, only the particular chain/sub-unit containing both XL peptides was chosen.

## RESULTS

### Protein structures can be automatically retrieved from AlphaFold

Our primary goal was to create an automated and easy-to-use tool to facilitate analysis of large-scale or proteome-wide XL-MS datasets. To this end, as a first step the three-dimensional structures of all proteins identified as having intra-protein crosslinks in each XL-MS experiment (as listed by UniProt ID in the relevant input file, see Experimental Methods) are automatically retrieved from the AlphaFold database **(Fig. 1)**. Importantly, the proteome database used for retrieving the structures from AlphaFold should be the same used for XL-MS data analysis, to ensure the same alignment of amino acid positions during mapping of the cross-linked peptides on to the 3D structures of the proteins.

### XL peptides can be automatically mapped to protein structures at proteome scale

To demonstrate the implementation of AlphaCross-XL, we first used our previously published XL-MS data datasets, obtained using two different cross-linkers (DSSO and PhoX) on surface-enriched proteins from the acute myeloid leukemia cell line Nomo-1^1^. The initial input into AlphaCross-XL is the output file from the in-house software xl-Tools used to first identify crosslinks present in this sample. In total, 138 proteins were identified to have crosslinks in this dataset and were included in the AlphaCross-XL analysis (**Supplementary Table 1**). As outputs of the AlphaCross-XL software, **Fig. 2A** shows a histogram of identified crosslinks length that, when mapped onto the compendium of AlphaFold structures, either satisfy or violate a user-defined C_α_-C_α_ distance that is expected to be the maximum such distance for the experimentally incorporated crosslinker molecule (here defined as 25 Å for crosslinker molecule DSSO). Notably, the AlphaCross-XL output data is as expected, where the large majority (89.6%) of detected crosslinks fall within the length threshold. **Fig. 2B** shows a similar plot though here recorded on a crosslink-by-crosslink basis. **Fig. 2C** shows an example PyMOL visualization of the mapped crosslinks on the AlphaFold-predicted structure of integrin α_L_, where multiple crosslinks are identified and all fall within the defined C_α_-C_α_ distance. Notably, as a straightforward visual cue, distance “satisfying” crosslinks are colored green throughout, whereas distance “violating” are in red. Such PyMOL structures are generated for every protein included in the analysis. We anticipate these initial output plots from AlphaCross-XL will specifically assist in structural refinement of predicted structures, or, alternatively, discovery of potentially new biology related to protein conformations separate from the minimal energy state predicted by AlphaFold, across a complex proteomic dataset.

**Figure 2:**
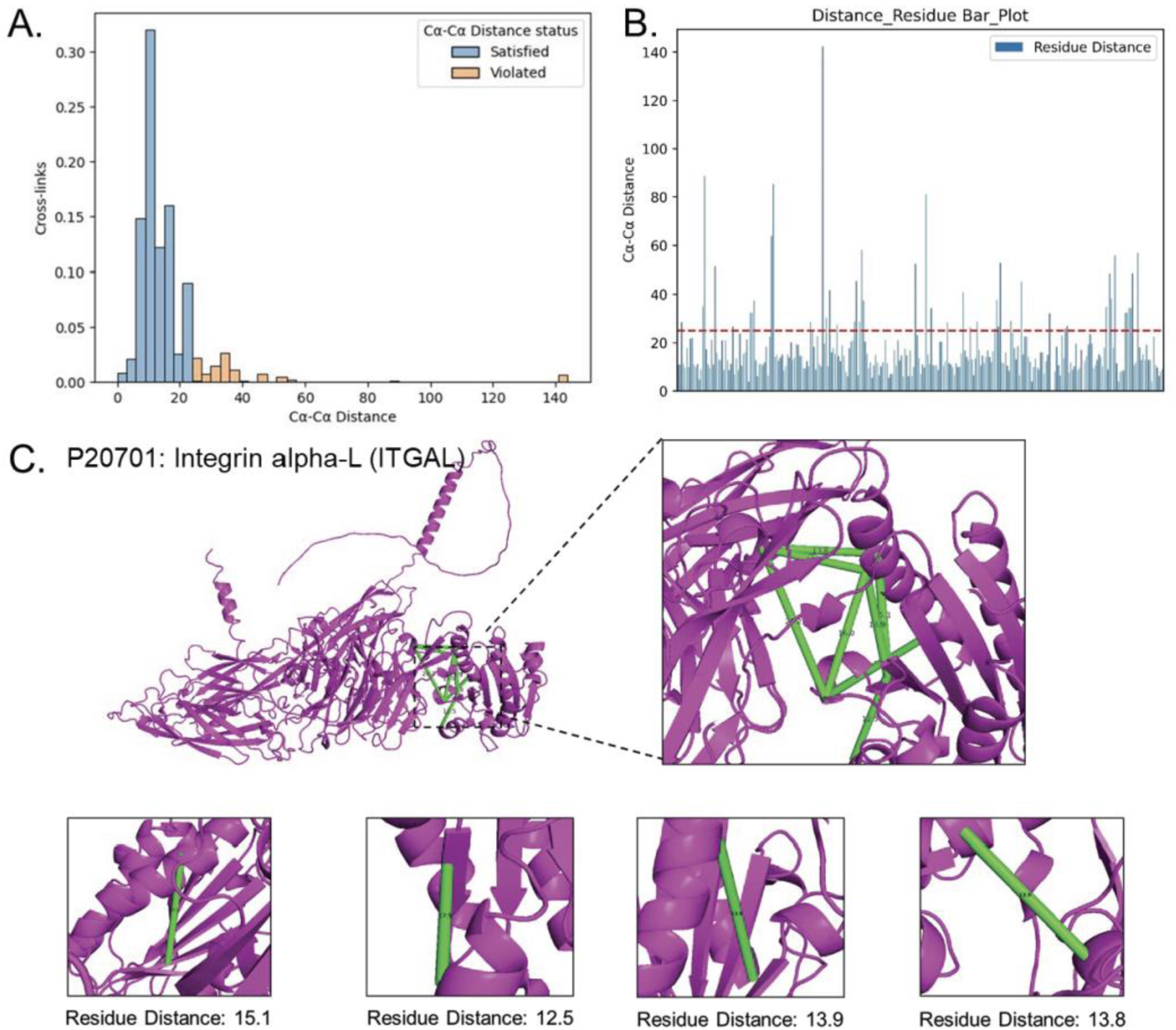
AlphaCross-XL output from the re-analysis of published structural surfaceomics data of AML models. **A)** Distribution plot of the proportion of residues against the residual distance, calculated based on the Euclidian distance between the target amino acid alpha-Carbon (set as Lysine for this dataset), where the individual bars are color coded based on the user-defined maximum distance, with crosslinks that satisfy the distance in blue and those that violate the distance in orange. **B)** Bar plot showing the crosslink distances with a red dashed horizontal line at the distance maximum for reference. **C)** Validation of crosslinks mapped onto ITGAL three-dimensional structure retrieved from AlphaFold, showing agreement between the structure and experimentally determined crosslinks.

Having started with the baseline AlphaFold structure for each protein, AlphaCross-XL also incorporates the confidence level of the mapped crosslinks in terms of the per-residue model confidence score (pLDDT) obtained from AlphaFold^18^. We believe this function will be important to help users further curate crosslinks of interest for downstream study; for example, “violating” crosslinks between residues of high confidence pLDDT may be considered more likely to reflect an accurate C_α_-C_α_ distance and therefore a true experimental deviation from the AlphaFold-predicted structure.

### XL peptide based structural inferences derived from AlphaFold are consistent with PDB protein structures

There may be cases where orthogonal validation and visualization of mapped crosslinks onto an experimentally obtained structure may be desired, as a complement to AlphaFold-predicted structures. To accommodate this need, AlphaCross-XL also allows mapping of the relevant XL peptides onto user-specified structures, including those obtained from the PDB. However, it is important to note that while AlphaFold-predicted structures will incorporate full-length, monomeric proteins, and our software is optimized for this type of structure, PDB structures will often include protein truncation variants (i.e. truncations amenable to recombinant expression of soluble protein) and/or multimeric or multi-chain structures. Particularly in truncated forms, we found the computational alignment of amino acid positions in an available PDB structure in comparison to an annotated UniProt sequence of the full-length protein could be challenging to consistently deconvolute. Despite these limitations, to demonstrate the robustness of our tool, as an initial test we manually curated available structures in the PDB to identify ten full-length (i.e. non-truncated), monomeric proteins (**Table 1**), which are anticipated to provide a more direct comparison to AlphaFold-predicted structures. Encouragingly, overall, we found high agreement between the output crosslink distances and visualization in this analysis, with 9 of 10 analyzed crosslinks concurring within 1 Å of the mapped C_α_-C_α_ distance based on either the AlphaFold-predicted structure or the PDB-determined structure (**Table 1**, **Fig. 3, Supplementary Fig. 2A-F**). The one larger deviation occurred in a structure of Voltage-dependent anion-selective channel protein 1 (VDAC1) (Uniprot: P21796) (**Table 1**). In this case, in AlphaFold both crosslinked peptides demonstrated high confidence structural predictions based on pLDDT scores (97.76 and 96.55, respectively). However, the comparator crystal structure from PDB was relatively low resolution (4.1 Å; PDB: 2JK4). Therefore, it is possible that the AlphaFold-mapped crosslink distance (13.73 Å) is in fact more reliable than that mapped from the PDB structure (10.91 Å) (**Supplementary Fig. 2C**). Taken together, these results further increase confidence in the use of AlphaFold predictions as the baseline structures for crosslink mapping in our tool. In this context, it is also important to note that using AlphaFold as a starting point enables us to evaluate crosslink spatial arrangements across a much broader swath of the proteome than the small portion experimentally available in the PDB.

**Figure 3:**
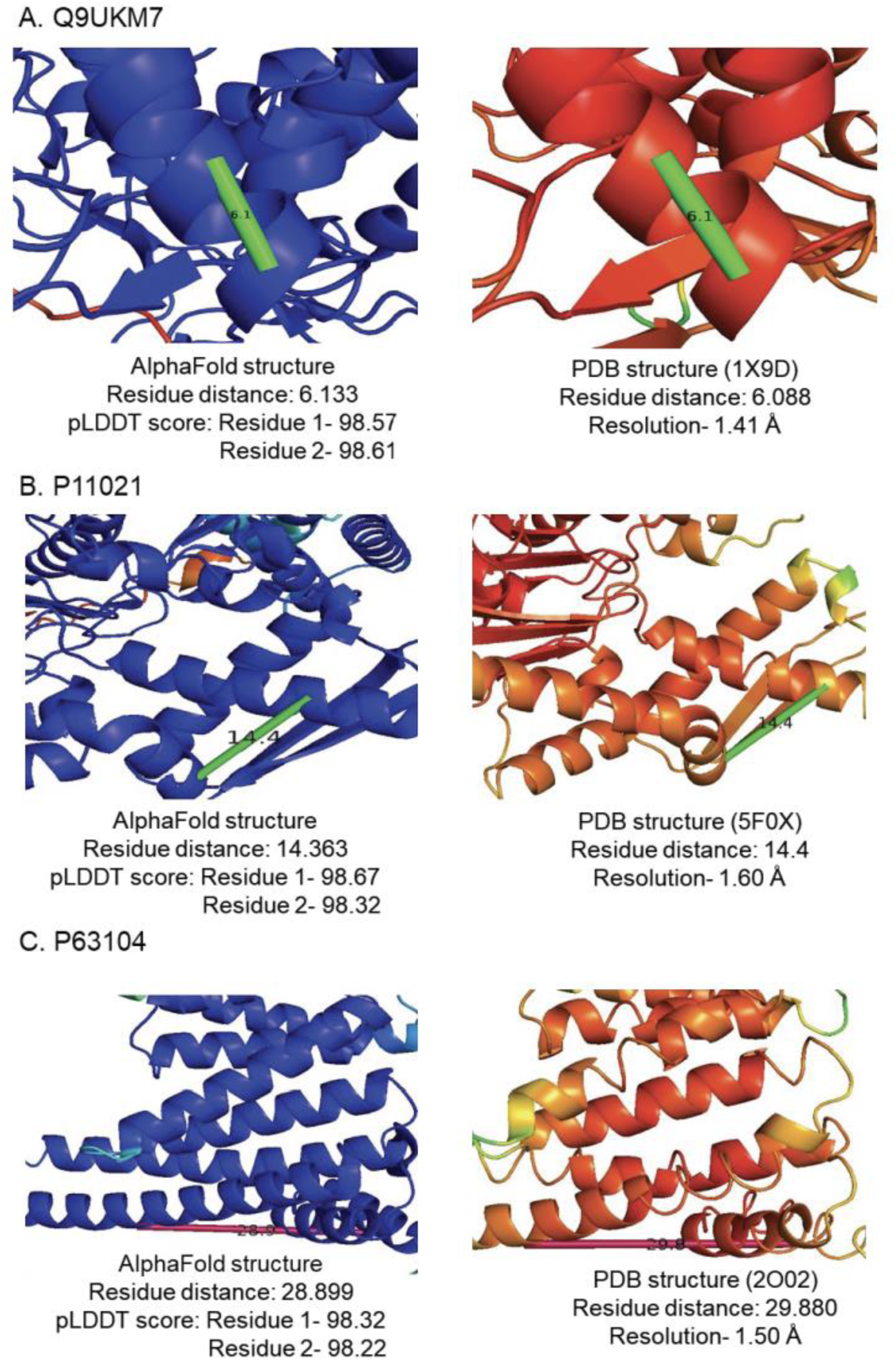
Comparison of the calculated residue distance in structures fetched from AlphaFold Protein Structure Database vs PDB. The residue distance from AlphaFold derived structures with residue pLDDT scores of 98.2 or greater were compared to residue distances calculated from deposited PDB structures with a resolution of 1.4 Å or greater for three proteins: **A)** MA1B1 (UniProt ID: Q9UKM7, PDB: 1X9D). **B)** BIP (UniProt ID: P11021, PDB: 5F0X), and **C)** 1433Z (UniProt ID: P63104, PDB: 2O02).

**Table 1:**
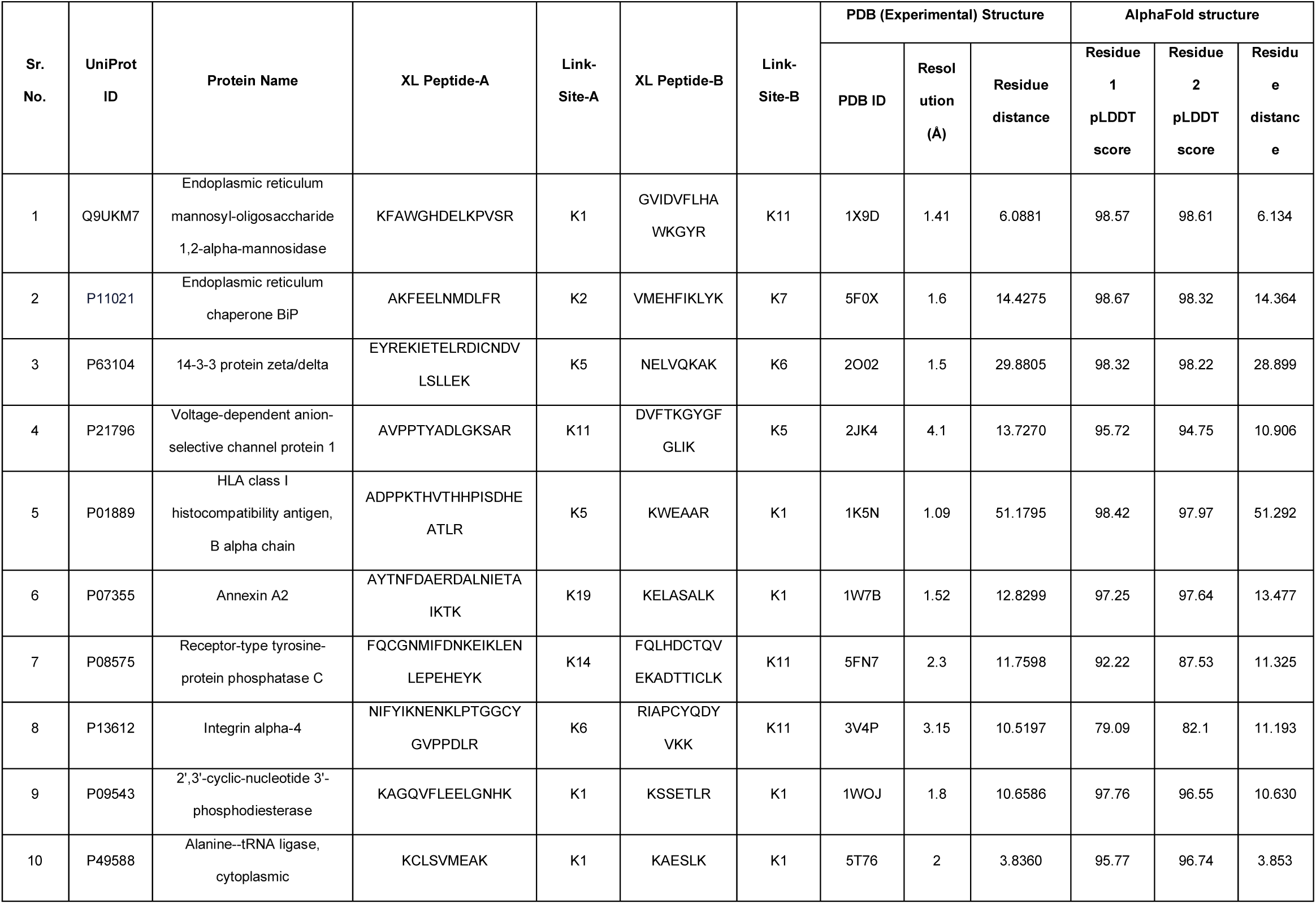
Comparison between the calculated distance in the AlphaFold-predicted and curated PDB-derived experimental structures.

### Validation of AlphaCross-XL workflow in myeloma and acute myeloid leukemia models

Another important test of the performance of our software is the consistency of generating comparative data across multiple related datasets. We therefore compared results from our XL-MS data derived from an AML cell line, as used in analyses in **Fig. 2**, to that of an independent XL-MS dataset from a multiple myeloma cell line, a different type of blood cancer (PXD059495). Using our data analysis pipeline, we sought to characterize and summarize the common crosslinked peptides detected in both the AML and myeloma datasets, which were both crosslinked using PhoX reagent (**Fig. 4A**). For display purposes, we show two examples in **Fig. 4**: detected crosslinks suggest that the protein STIM1 (UniProt ID: P01893) contains crosslinks that violate the distance constraints (shown in red, **Fig. 4B**), whereas TECR (UniProt ID: Q9NZ01) adopts the AlphaFold-predicted structure, as all identified crosslinks satisfy the distance constraints, shown in green (**Fig 4C**). In total, we identified 31 overlapping crosslinks on 28 different proteins, with only 10% (3/31) of the identified overlapping crosslinks violated the distance constraints of PhoX (**Supplementary Table 2**). Taken together, this high degree of overlap obtained from two independent datasets further suggests that AlphaCross-XL can provide reproducible the C_α_-C_α_ distances that can be rapidly visualized for purposes of data analysis and crosslink curation.

**Figure 4:**
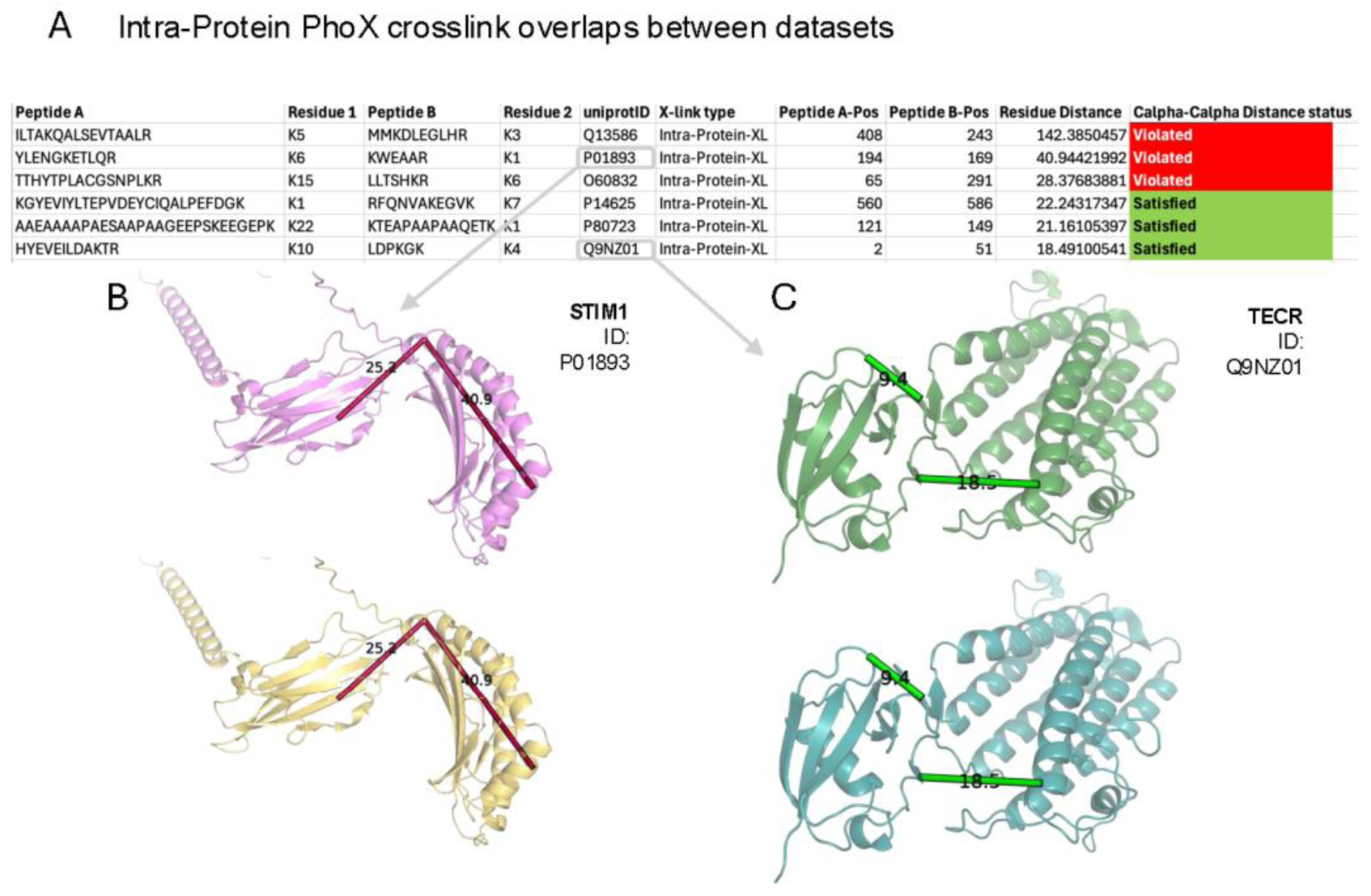
Overlap of intra-protein crosslinks from two independent PhoX labeled structural surfaceomics datasets. **A)** Table of common crosslinks across the datasets. **B-C)** Crosslinks that violate and satisfy the distance constraints respectively, identified in both datasets, as shown here of the protein STIM1 (UniProt ID: P01893) and TECR (UniProt ID: Q9NZ01).

### Validation of AlphaCross-XL workflow in published XL-MS datasets

We further validated AlphaCross-XL on published datasets from *E. coli* to demonstrate the utility of our tool in bacterial proteomes. We analyzed data from Ruwolt *et al*., which contained PhoX crosslinked *E. coli*^19^. Crosslinks were identified with Proteome Discoverer 3.0 software (Thermo Fisher Scientific) with Xlinkx node 3.0, and the CSV output of the search software was used as the input for AlphaCross-XL. Using our software, we were able to generate structures with informative crosslinks that demonstrate agreement between the experimentally determined and AlphaFold predicted structures (**Supplementary Fig. 3**). Through AlphaCross-XL, we mapped a total of 480 crosslinks onto AlphaFold-predicted proteins; 125 of the mapped proteins contained three or more high confidence crosslinks.

## DISCUSSION

XL-MS driven analysis is emerging as an important approach in structural biology^12,25,26^. Unlike conventional biophysical methods using recombinant proteins, the ability of XL-MS to provide structural information of proteins functioning in their native biological context in real-time may elucidate more physiologically-relevant protein structures. One such application has recently been reported by our group, where we developed a technology called “structural surfaceomics” by integrating XL-MS with surface protein biotinylation via cell surface capture^1^. This approach enables unbiased identification of cancer-specific cell surface protein conformations that can then be targeted with novel therapies. Our prior study, however, was limited in throughput as it involved manual mapping of XL peptides onto protein structures obtained from the PDB. This prior approach is not only incredibly labor-intensive for proteome-wide XL-MS analyses, but also is constrained by the number of available structures in the PDB, which are particularly limited for membrane proteins. These hurdles in proteome-wide XL-MS analysis and crosslink visualization, also faced by others^2^, inspired us to develop AlphaCross-XL. This software tool streamlines this curation and filtration of crosslinks of potential biological interest through automated mapping of crosslinks onto 3D protein structures derived from AlphaFold. We designed AlphaCross-XL to be applicable to any type of XL-MS study. Alterations in protein conformation are general phenomena in biology. The ability to rapidly detect such alternative protein conformations in complex systems could strengthen our ability to address a range of biological questions including those in cancer, infectious diseases, neurological disorders, and others.

We believe AlphaCross-XL is a first-in-class tool enabling proteome scale structural mapping of crosslinked peptides obtained from XL-MS experiments. Though XL-MS experiments are typically based on lysine targeted crosslinking using NHS ester chemistry of the crosslinkers^4^, our tool allows for visualization of XL-MS data generated with crosslinkers targeted to other amino acids, including multi-amino acid targeted promiscuous crosslinkers, such as the crosslinker succinimidyl diazirine carbamate (dizSEC) and succinimidyl diazirine sulfoxide (SDASO)^7,8^. AlphaCross-XL is user friendly with an intuitive GUI for setting analysis parameters, and, in addition to readily interpretable output plots of crosslink distance (**Fig. 2**), the software also outputs a separate .PSE PyMOL session file for each crosslink located in an output directory of the protein labeled by UniProt ID upon completion of analysis, to facilitate visualization of XL peptides. This visualization also enables easy color code-based discrimination of satisfied and violated crosslinks based a user-defined distance threshold. Incorporation of pLDDT scores into crosslink mapping also assists in the triage of the most biologically relevant crosslinks for downstream investigation. We do note that our tool does not create scoring or other complex functions for the generation of crosslink guided structures, such as the recently reported tool AlphaLink^27^ and analysis pipeline reported by Manalastas-Cantas *et al*.^28^. In contrast, the primary focus of our tool is a rapid search and interpretation software for high throughput crosslink structural mapping.

Currently, our tool supports AlphaFold for seamless structural mapping of XL peptides at proteome scale, eliminating the restriction of mapping crosslinks onto deposited PDB structures. Several groups have worked towards XL-MS data visualization, such as xiVIEW, which utilizes NGL to display 3D protein structures with mapped crosslinks and allows users to export into PyMOL^29^. However, this tool requires a PDB file, either from the official repository or created locally, which is an additional input parameter the user needs to consider. Other tools with PyMOL integrations exist such as PyXlinkViewer^30^ and XlinkCyNET^31^, however these do not support AlphaFold and they also require a manually-inputted PDB file.

Notably, though, our tool can still be applied to PDB structures, if desired, and, as we showed (**Table 1**), the concordance between crosslinks mapped to either AlphaFold and PDB-based structures is strong in the context of high-resolution X-ray data. Additionally, our tool can also support other structural databases, such as cryoEM structures from the Electron Microscopy Data bank (EMDB). Notably, modeling integrated structures from XL-MS and cryoEM is currently a rapidly advancing field^32,33^.

AlphaCross-XL has been successfully implemented to structural mapping and visualization of XL-MS search results from various XL-MS analysis software suites and search engines such as pLink 2.0^14^, Kojak^15^, SCOUT^16^, and XiSearch^17^. As a limitation, the current version of AlphaCross-XL supports only intra-protein crosslinks for making structural inferences. Future versions will also ideally support analysis of inter-protein crosslinks, facilitating an even broader scope of protein complex modelling and protein-protein interaction mapping in proteome-wide analyses.

## Author Contributions

KM, SS, APW, DB, SVS, AS, AZ, and AK have conceptualized the software features and verified software integrity. SVS, AZ, and DB have designed and developed the software. KM, AK, YH, and RV have generated the samples, and the input files used for analysis and testing of the software. SVS, DB and AV have completed the visualization. IA has contributed to the software development and software documentation. SVS, DB, KM, AV, AK, SS and APW wrote the manuscript and obtained funding for the project. All authors contributed to the article and approved the submitted version.

## Code Availability

Web version of AlphaCross-XL is available at https://tinyurl.com/alphacrossxl-202501

Instruction Manual, Tutorial Video and Files for testing purposes, and local version of AlphaCross-XL are available at https://github.com/sanjyotshenoy/alphacross-xl

## Acknowledgements

Authors would like to acknowledge Abhilash Barpanda for providing his inputs in this work.

## Supplemental Data

This article contains supplemental data^1,31^.

## ABBREVIATIONS

XL-MS: crosslinking mass spectrometry
XL peptides: crosslinked peptides
3D: three-dimensional
PDB: Protein Data Bank
pLDDT: predicted local distance difference test
CAR-T: chimeric antigen receptor T-cells
CSV: comma-separated values
GZIP: GNU zip
FASTA: Fast Adaptive Shrinkage Threshold Algorithm
CIF: Crystallographic Information File
PSE: PyMOL Session

