## Supplemental Data for "AlphaCross-XL: a seamless tool for automated and proteome-scale mapping of crosslinked peptides onto three-dimensional protein structures"

**List of Materials**

**Table S1 (attached as separate file).** Detailed Summary of the DSSO based XL-MS data. This spreadsheet is the processed data file from Mandal *et al.,* 2023[1], containing the identified crosslinks from DSSO-linked cell surfaceomics of the AML cell line, Nomo-1. This spreadsheet was directly uploaded to AlphaCross-XL in csv format.

**Table S2 (attached as separate file).** Comparison of common crosslinks from two independent PhoX XL-MS (structural surfaceomics) datasets. This spreadsheet lists the common crosslinks between two independent PhoX-linked cell surfaceomics datasets in different hematologic malignancy cell lines. The datasets compared were of the AML cell line Nomo-1, taken from Mandal *et al.* 2023 and of the myeloma line AMO-1 (PXD059495).

**Table S3 (attached as separate file).** This spreadsheet is the processed data file from Ruwolt *et al.,* 2023[2], containing the identified crosslinks from PhoX-linked bacterial lysates. This spreadsheet was directly uploaded to AlphaCross-XL in csv format.

**Figure S1.** The AlphaCross-XL GUI and its input requirements.

**Figure S2.** Extended comparison of the calculated residue distance in structures fetched from AlphaFold Protein Structure Database vs PDB.

**Figure S3.** AlphaCross-XL utility in crosslinked bacterial proteomes.

**Supplementary References**

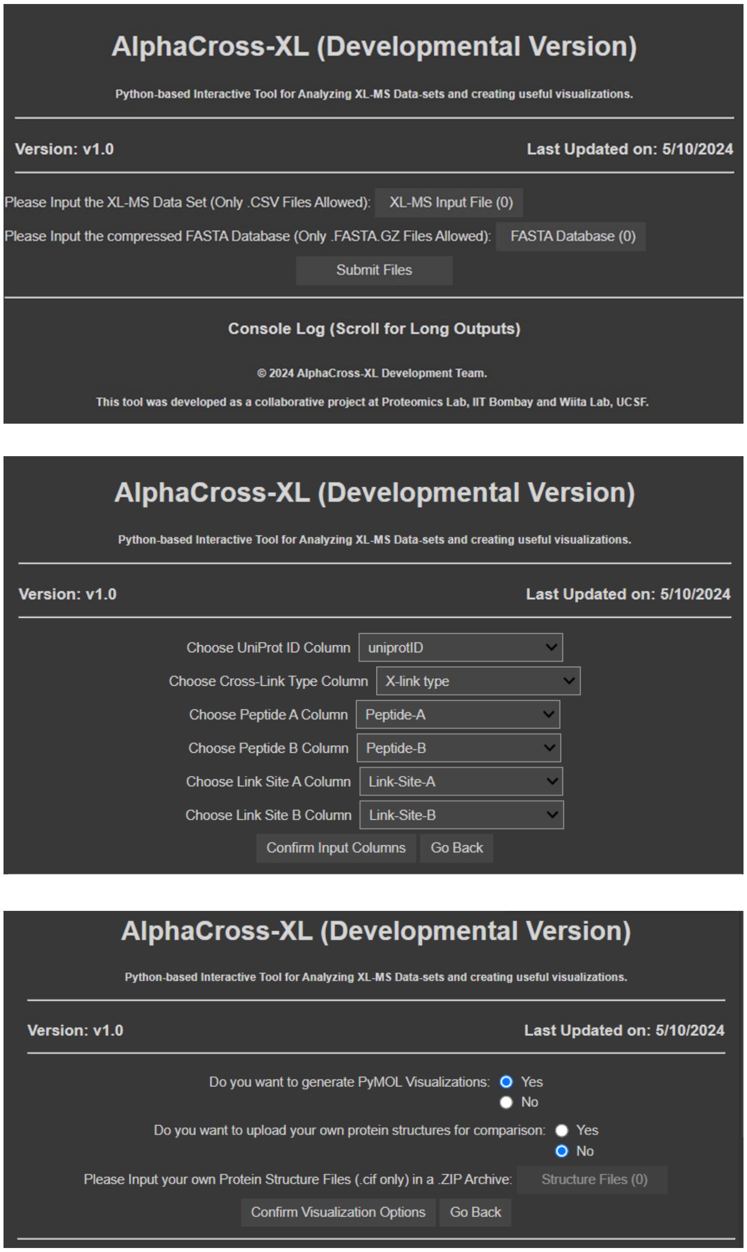


**Figure S1: The AlphaCross-XL GUI and its input requirements. A)** Represents the homepage interface of AlphaCross-XL. The initial steps include choosing input files, FASTA sequence database and parameters required for the AlphaCross-XL analysis (reviewed) file along with the threshold. **B)** Shows the following step of selecting headers for the analysis which includes UniProt ID, Cross-Link Type, Peptide Sequences and peptide site specifications. **C)** Deciphers the PDB comparison analysis interface where a custom database of curated PDB structures could be uploaded by the user to get a comparison between PDB and AlphaFold structure-based distance calculation.

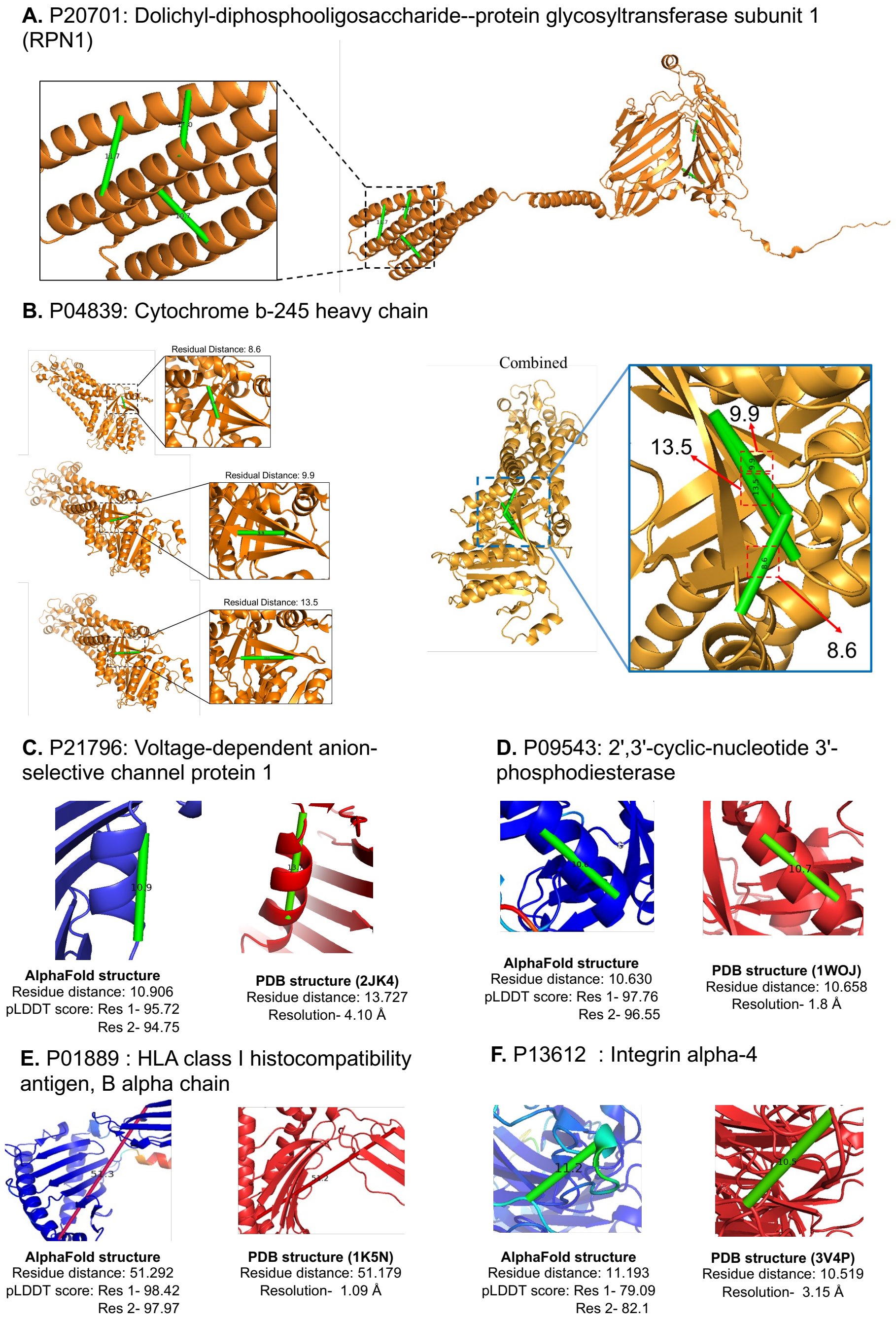


**Figure S2: Extended comparison of the calculated residue distance in structures fetched from AlphaFold Protein Structure Database vs PDB. A-B)** Cross-links available for multiple residues for RPN1 and CY24B heavy chain. **C-F)** Comparison between the calculated residue distance fetched from AlphaFold Structure and PDB structure for 4 proteins which includes: **C)** VDAC 1. **D)** CN37. **E)** HLAB. **F)** ITA4.

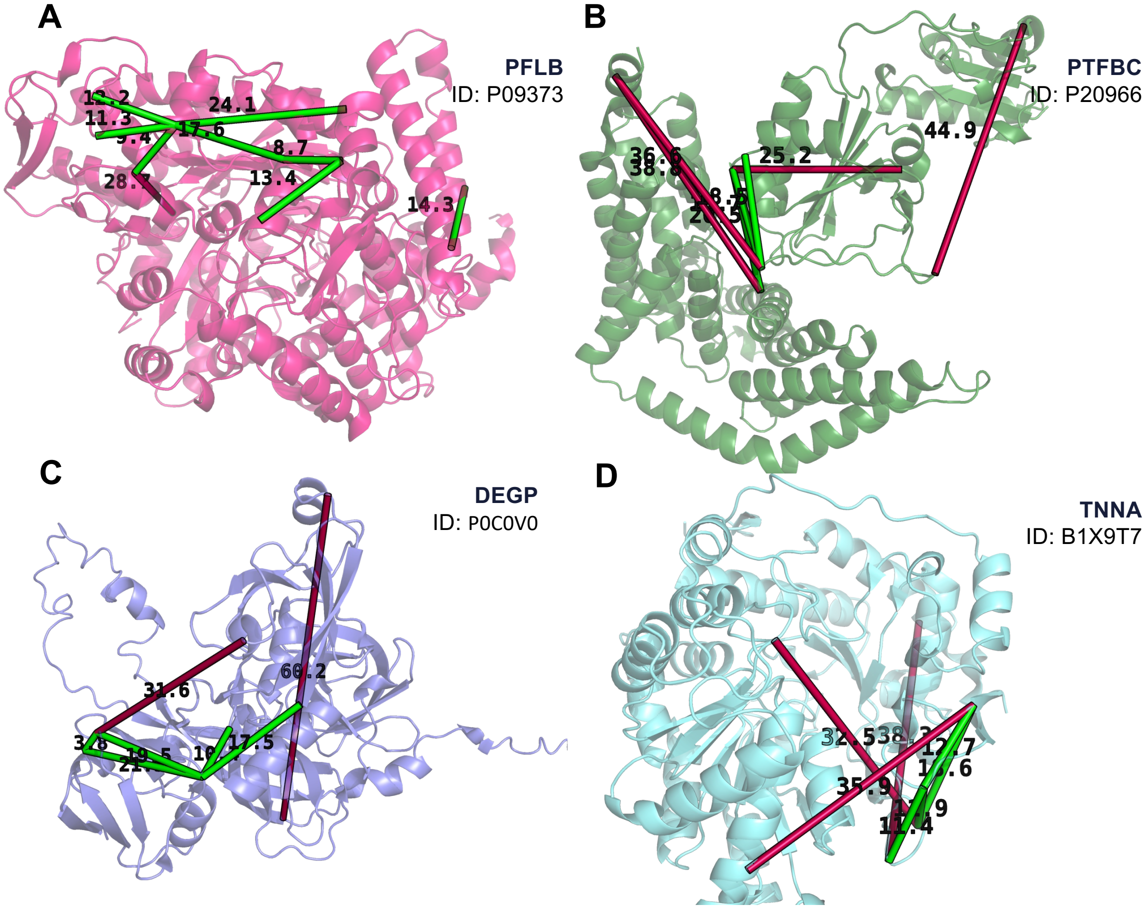


**Figure S3: AlphaCross-XL utility in crosslinked bacterial proteomes. A-D)** Output of Alpha-Cross-XL from PhoX crosslinked *E. coli* lysates. XL-MS dataset from Ruwolt *et al.,* 2023[2]. Crosslinks for the following proteins are shown: **A)** PFLB. **B)** PTFBC. **C)** DEGP. **D)** TNNA.
